# Hippocampal hyperactivity in a rat model of Alzheimer’s disease

**DOI:** 10.1101/2020.06.09.141598

**Authors:** Liudmila Sosulina, Manuel Mittag, Hans-Rüdiger Geis, Kerstin Hoffmann, Igor Klyubin, Yingjie Qi, Julia Steffen, Detlef Friedrichs, Niklas Henneberg, Falko Fuhrmann, Daniel Justus, Kevin Keppler, A. Claudio Cuello, Michael J. Rowan, Martin Fuhrmann, Stefan Remy

**Affiliations:** Neuronal Networks Group, German Center for Neurodegenerative Diseases (DZNE), Bonn, Germany; Neuroimmunology and Imaging Group, German Center for Neurodegenerative Diseases (DZNE), Bonn, Germany; Light Microscopy Facility, German Center for Neurodegenerative Diseases (DZNE), Bonn, Germany; Department of Cellular Neuroscience, Leibniz Institute for Neurobiology, Magdeburg, Germany; Department of Pharmacology and Therapeutics, Watts Building, Trinity College, Dublin 2, Ireland; Department of Pharmacology and Therapeutics, McGill University, 3655 Sir-William-Osler Promenade, Room 1210, Montreal, QC H3G1Y6, Canada

## Abstract

Neuronal network dysfunction is a hallmark of Alzheimer’s disease (AD). However, the underlying pathomechanisms remain unknown. We analyzed the hippocampal micronetwork in a rat model of AD at an early disease stage at the beginning of extracellular amyloid beta (Aβ) deposition. We established two-photon Ca^2+^-imaging *in vivo* in the hippocampus of rats and found hyperactivity of CA1 neurons. Patch-clamp recordings in brain slices *in vitro* revealed changes in the passive properties and intrinsic excitability of CA1 pyramidal neurons. Furthermore, we observed increased neuronal input resistance and prolonged action potential width in CA1 pyramidal neurons. Surprisingly, all parameters measured to quantify synaptic inhibition and excitation onto CA1 pyramidal neurons were intact suggesting a cell immanent deficit. Our data support the view that altered intrinsic excitability of CA1 neurons may precede inhibitory dysfunction at an early stage of disease progression.

## Introduction

Defining the earliest neuronal changes on a cellular and network level of brain organization that precede the clinical manifestation of Alzheimer’s disease (AD) remains an important task with potential relevance for early diagnosis, treatment and prevention. The majority of previous studies that have addressed changes of neuronal integrity and excitability were performed at disease stages, where amyloid beta (Aβ) plaques were reliably detected within the brain region of interest^1, 2^. These studies report changes in the intrinsic excitability, the morphology and the *in vivo* activity of neurons in the presence of amyloid beta^2^. Common pathophysiological motifs on the single cell level were structural alterations such as reduced dendritic arborization, spine loss, synapse loss and reduced cell size^1, 3, 4, 5, 6, 7^, which were observed in conjunction with higher neuronal excitability due to altered passive and active neuronal electrical properties^8^. On the circuit level reduced synaptic inhibition has been identified in several rodent models of beta amyloidosis. Inhibitory dysfunction has been associated with the occurrence of aberrant activity patterns such as neuronal hyperactivity^9^, hypersynchronization within neuronal networks^10, 11, 12^ and impaired neuronal oscillations^13, 14^. A current view is that these aberrant forms of neuronal activity are mediated by the action of soluble forms of Aβ, which shift the balance of excitation and inhibition towards excitation^15^. Evidence supporting this view are the prevention of neuronal hyperactivity by □-secretase inhibitors^16, 17^ and the restoration of physiological activity and oscillatory patterns as well as the amelioration of memory deficits by pharmacological or optogenetic enhancement of inhibition^18, 19, 20, 21^. Among the studies that addressed the earliest changes of cellular function at pre-plaque stages the majority report alterations on the synaptic level including synaptic plasticity^22, 23, 24^, but evidence for neuronal hyperactivity within the hippocampus has also been provided^16, 25, 26, 27^. Here, we investigated whether intrinsic neuronal excitability changes and inhibitory circuit dysfunction are mechanistically involved in neuronal hyperactivity within the CA1 hippocampal subfield at an early stage of extracellular Aβ deposition. We used transgenic McGill-R-Thy1-APP rats (APPtg), a model with AD-like pathology overexpressing hAPP (human amyloid precursor protein) that exhibits a slow progression of amyloidosis, synaptic and memory deficits^23, 28^.

## Results

### Increased Ca^2+^-transient frequency of hippocampal CA1 pyramidal neurons

Hyperactivity of neurons in the cortex and hippocampus of mice has been previously shown to occur in proximity of Aβ-plaque depositions^2^ and at pre-plaque stage^16^. Therefore, we first addressed, whether hyperactive neurons are equally present at an early stage of Aβ-plaque deposition in the hippocampus of our transgenic rat model. We used homozygous Thy1- APP-tg (APPtg) and non-transgenic wildtype (WT) rats. We established a cranial hippocampal window that enabled us to carry out two-photon *in vivo* imaging in rats (Fig. S1). We built a headpiece that was permanently attached to the skull with dental acrylic (Fig. S1a). A headbar served as temporary adapter to attach the rat in anesthesia to the *in vivo* headpost stand (Fig. S1b, c; see Material and Methods). Since microgliosis and reactive astrocytes have been shown to be associated with hippocampal window implantation^29^, we investigated both parameters (Fig. S1d). We found that the number of microglia were not different between the implanted and non-implanted hemispheres of APPtg and WT rats (Fig. S1e, f). While the number of astrocytes were decreased on the ipsilateral side in WT rats, the number of astrocytes was comparable between ipsi- and contralateral hemisphere of APPtg rats and the contralateral hemisphere of WT rats (Fig. S1g, h). These results indicated that inflammation on the ipsilateral side decayed to levels of the contralateral side at least 3 weeks after surgery in both APPtg and WT rats. This is in line with previous results obtained in mice^29^. Previous studies on our transgenic rat model found a start of Aβ-plaque deposition in the hippocampus at 6-9 months of age^23, 28, 30^. In order to confirm this result, we analyzed extracellular Aβ-plaques within our experimental groups at 6-9 months of age. Using an APP-specific (6E10), an Aß-specific (4G8) antibody and Thioflavin S staining, we only detected a few Aβ-plaques in the hippocampus of APPtg rats (Fig. S2). In contrast, Aβ-plaque load was strongly increased in APPtg rats older than 18 months of age Fig. S2f).

In order to assess if chronic hippocampal windows in the rat allowed for robust recording of Ca^2+^-transients in populations of CA1 neurons we injected a recombinant adeno-associated virus encoding the genetically encoded Ca^2+^-indicator GCaMP6m driven by a synapsin promoter (*rAAV-hSyn-GCaMP6m*). Next, we performed two-photon *in vivo* imaging of GCaMP6m expressing CA1 neurons in isoflurane (1%) anesthesia (Fig. 1a, b). APPtg rats exhibited significantly more Ca^2+^-transients per neuron than WT rats (Fig. 1b). In total we monitored 1623 neurons in WT rats (WT, 6-9 months old, 31 imaging areas in 5 animals) and 2051 neurons in APPtg rats (APPtg, 6-9 months old, 35 imaging areas in 6 animals). We monitored the occurrence of spontaneous somatic Ca^2+^-transients as an indicator of neuronal activity. First, we detected a prominent shift in CA1 neurons towards higher frequencies of Ca^2+^-events in APPtg compared to WT rats (Fig. 1c). We further assessed CA1 network activity by determining the proportion of active neurons that exhibited at least one Ca^2+^- transient throughout the two-minutes recording (Fig. 1d). Furthermore, we calculated the mean Ca^2+^-transient frequency (Fig. 1e). We found that the proportion of active CA1 neurons and their mean Ca^2+^-transient frequency was significantly increased in APPtg compared with WT rats (APPtg: 69.63 [56.06; 80.21] % vs. WT: 30.21 [7.83; 55.26] %; APPtg: 6.51 [2.46; 8.41] min^-1^ vs. WT: 0.60 [0.14; 1.41] min^-1^). In order to further specify this finding, we classified the cells into categories of silent, moderately active and hyperactive neurons according to previous reports^2, 16^. In WT rats (Fig. 1f) the majority (>95%) of CA1 neurons were either categorized as silent (61%) or moderately active (36%). Hyperactive neurons only made up for a marginal proportion (3%). On the other hand, in APPtg rats the proportions were strongly shifted (Fig. 1g). Hyperactive CA1 neurons formed the largest population (38%), exceeding the moderately active population (34%) and the silent population (28%). Comparing the proportions of moderately active and hyperactive CA1 neurons between WT and APPtg rats revealed no change in the population with moderately active neurons (Fig. 1h; WT: 8.72 [7.48; 33.79] % vs. APPtg: 34.49 [19.16; 42.59] %). On the other hand, the proportion of hyperactive CA1 neurons was significantly increased in APPtg compared to WT rats (Fig. 1i; APPtg: 34.53 [16.29; 51.71] % vs. WT: 0.92 [0; 11.66]). These results show an increased fraction of hyperactive neurons in dorsal CA1 of APPtg rats. In order to determine whether an increase in glutamatergic synaptic transmission contributed to the observed CA1 pyramidal neuron hyperactivity, we measured hippocampal electrically evoked synaptic input-output relationships in a freely-behaving rats. Similar to our findings in younger animals^23^, baseline transmission at CA3 to CA1 synapses was statistically indistinguishable between 6–9 months old WT and APPtg rats (Fig. 1k). These results do not support a role for increased glutamatergic synaptic transmission as a cause of the neuronal hyperactivity in the CA1 area of APPtg rats.

**Figure 1.**
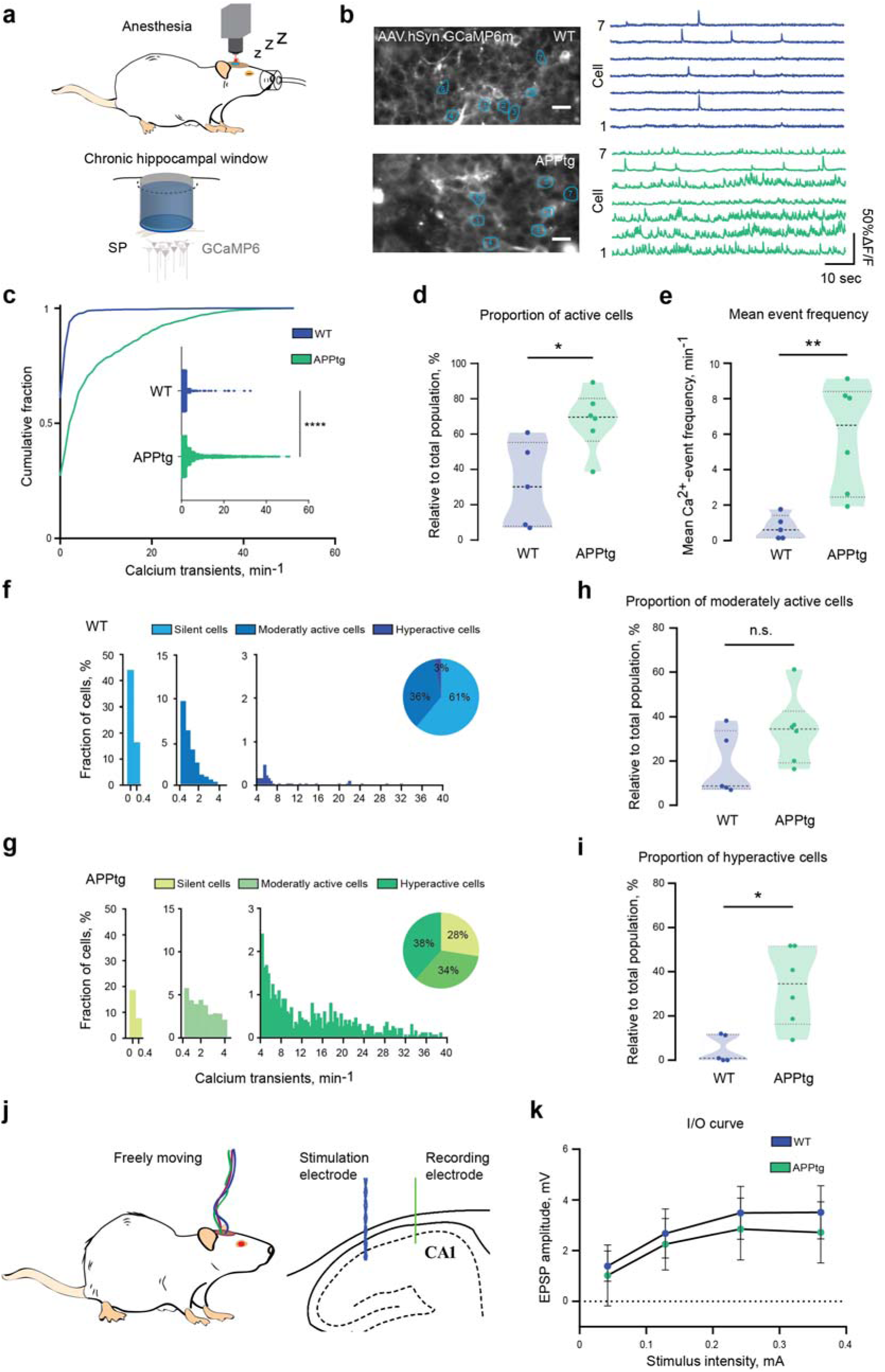
Imaging of CA1 neurons in the dorsal hippocampus of APPtg rats. (a) Schematic illustration of a chronic cranial window in rats that allows imaging of neurons in stratum pyramidale (SP) in the CA1 region of the dorsal hippocampus. (b) Exemplary picture of CA1 neurons and respective calcium traces in WT and APPtg rats. (c) Cumulative density plot of pooled cells of WT (n=1623) and of APPtg (n=2037) comparing calcium event frequencies (Kolmogorov Smirnov test, **** p<0.0001, D=0.468). (d) Comparison of Proportion of active cells (cells that show at least one Ca^2+^-transient during recording time) in WT rats (n=5, 30.21 [7.83; 55.26] %) and APPtg rats (n=6, 69.63 [56.06; 80.21] %) averaged over animals (Mann Whitney test, p=0.017). Values are displayed as median [25% Percentile; 75% Percentile]. (e) Comparison of mean Ca^2+^-transient frequency in WT rats (n=5, 0.60 [0.14; 1.41] min^-1^) and APPtg rats (n=6, 6.51 [2.46; 8.41] min^-1^) averaged over rats (Mann Whitney test, p=0.004). (f) Pooled cells from all WT rats categorized into silent cells (<0.4 Ca^2+^-transients/min), moderately active cells (0.4-4 Ca^2+^-transients/min) and hyperactive cells (>4 Ca^2+^-transients/min). (g) Pooled cells from all APPtg rats categorized into silent cells, moderately active cells and hyperactive cells. (h) Comparison of moderately active cells (0.4-4 Ca^2+^-transients/min) in WT rats (n=5, 8.72 [7.48; 33.79] %) and APPtg rats (n=6, 34.49 [19.16; 42.59] %) averaged over animals (Mann Whitney test, p=0.178). (i) Comparison of hyperactive cells (>4 Ca^2+^-transients/min) in WT rats (n=5, 0.92 [0; 11.66] %) and APPtg rats (n=6, 34.53 [16.29; 51.71] %) averaged over animals (Mann Whitney test, p=0.017). (j) Schema of recoding (green) and stimulation (blue) electrode configuration in the area of CA1 of the dorsal hippocampus of freely behaving rat. (k) Baseline synaptic efficacy, as measured by the electrically evoked field EPSP input-output curve, was not different between WT and APPtg rats (Two-way RM ANOVA). Values are displayed as mean ± SD. Scale bars=20 *µ*m.

### Intact parameters of inhibitory neurotransmission in CA1 of APPtg rats

Previous studies in mouse models of AD revealed deficits on the level of synaptic inhibition that contributed to neuronal network dysfunction and were thought to underlie learning and memory impairments^10, 21, 31, 32^. Therefore, we tested whether APPtg rats also exhibited impaired inhibition^23, 28^. We carried out alveus stimulation in brain slices *in vitro* in order to recruit full recurrent inhibition and to further dissect the proximal and distal dendritic inhibitory component according to established paradigms^33, 34^ (Fig. 2). With increasing stimulation intensity, we did not observe differences between the input-output curves of WT and APPtg rats (Fig. 2a, b). In agreement with these finding, the maximal IPSP amplitude remained unchanged (WT: -4.57±1.39 mV; APPtg: -3.83 ± 1.49 mV, Fig. 2c). We then used 5Hz stimulation to investigate the frequency dependent attenuation of IPSPs within the range of physiologically observed theta oscillations (Fig. 2d). Moreover, detailed analysis of IPSP kinetics did not reveal differences neither in mean hyperpolarization, nor in IPSP rise and decay time constants between WT and APPtg rats (Fig. S3a-c). No difference in the frequency dependent dynamics of the IPSP amplitude was observed (Fig. 2f; percent reductions 10^th^ versus 1^st^ IPSP: WT 67.80 [54.08; 77.30] % vs. APPtg 69.29 [56.13; 73.82] %). Finally, we applied 100 Hz burst stimulation at theta frequency (5 Hz). This protocol allowed to distinguish the inhibitory circuitry targeting more distal dendrites from proximal inhibition, as a relative reduction of the first intra-burst response predominantly represents proximal inhibition whereas the 3^rd^ intraburst component represents the distal component^33, 34^ (Fig. 2g-h). First, we analyzed the compound IPSP resulting from three stimulations at 100 Hz burst frequency. The frequency dependent reduction in IPSP amplitude was unchanged comparing WT and APPtg rats (Fig. S3d) and reached previously described values of 60%^33^. Furthermore, we analyzed distal and proximal inhibition separately. Reduction of the first component at 100 Hz burst as well as the third component of first pulse in comparison to the last pulse did not deviate between WT and APPtg rats (Fig. 2i). These results suggest that neither proximal nor distal inhibition onto CA1 pyramidal neurons is affected in 6-9 months old APPtg rats.

**Figure 2.**
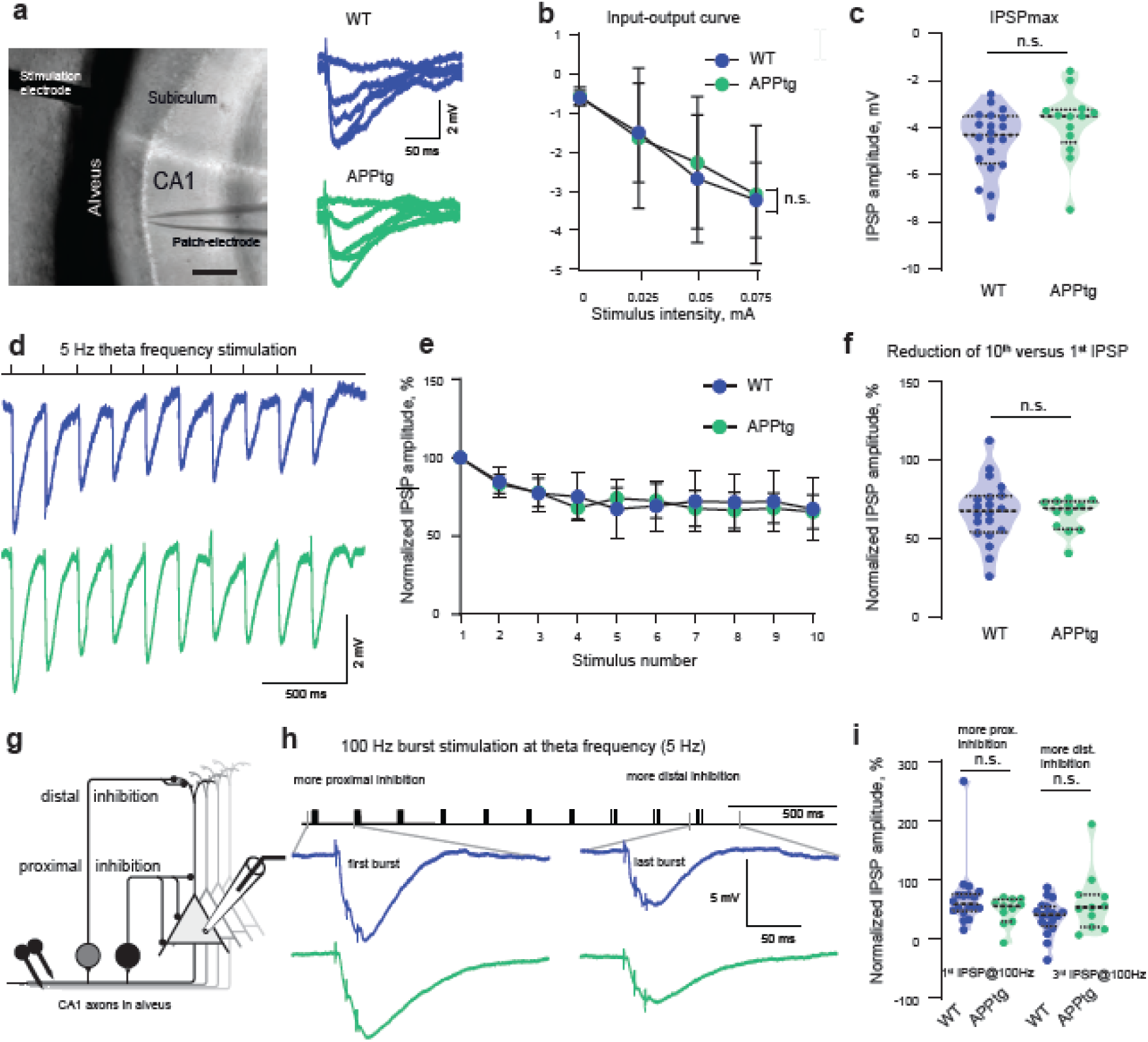
Intact proximal and distal inhibition of CA1 neurons in APPtg rats. (a) Experimental setup with stimulation electrode in alveus and patch pipette in CA1 area in rat hippocampal slice. Scale bar, 200 *µ*m. Representative examples of evoked IPSPs with stimulation intensity increasing from 0 mA to 0.1 mA in 0.025 mA step. Blue lines correspond to WT rats; green to APPtg rats. (b) Extracellular stimulation of a CA1 pyramidal neuron with increasing stimulation intensity of (0, 0.025, 0.05 and 0.075). Input-output relationships were not different between WT and APPtg rats (Two-way RM ANOVA, Sidak’s multiple comparisons test). (c) Maximal IPSP amplitudes were not different between WT (n=20 in 12 animals) and APPtg (n=13 in 7 animals) rats (unpaired t-test, p=0.128). (d) Representative examples of 5 Hz alveus stimulation in WT and APPtg CA1 pyramidal neurons. (e) Normalized IPSP amplitude was not significant different between WT (n=20 in 12 animals) and APPtg (n=12 in 6 animals) rats (Two-way RM ANOVA,Sidak’s multiple comparisons test). Error bars represent mean ± SD. (f) Reduction in IPSP amplitudes 10^th^ (last) vs. first stimulation (5 Hz), was not different between WT (n=20 in 12 animals) and APPtg (n=12 in 6 animals) (Mann-Whitney test, p=0.893). (g) Schematic illustration of alveus stimulation and two types of interneurons targeting more proximal (black) or more distal (grey) dendrites of CA1 pyramidal neuron. (h) Alveus 100 Hz burst stimulation at theta frequency (5 Hz) allows distinguishing proximal and distal inhibition. Lower traces: compound IPSPs in WT and APPtg rats in response to the first compared to the 10^th^ repeated burst. (i) Reduction in the first IPSP amplitudes at 100 Hz burst of the first vs. last stimulation burst (100 Hz burst at 5 Hz), corresponding to more proximal inhibition, was not different between WT (n=18 in 12 animals) and APPtg (n=11 in 6 animals) (Mann-Whitney test, p= 0.159). Reduction in the third IPSP amplitude at 100 Hz burst of the first vs. last burst, corresponding to more distal inhibition, was not different between WT (n=18 in 12 animals) and APPtg (n=11 in 6 animals) (Mann-Whitney test, p=0.238).

### Increased intrinsic excitability in APPtg rats

Since the measured inhibitory parameters were intact, we assumed that other mechanisms might be responsible for CA1 neuronal hyperactivity. We have previously shown by using computational modeling on reduced neuronal morphologies of APP/PS1 mice that decreased membrane area was associated with increased neuronal input resistance resulting in increased neuronal excitability^8^. We hypothesized that similar as in mice, rats might show increased input resistance. Therefore, we carried out electrophysiological recordings in 15 WT and 9 APPtg rats. We observed a difference in the resting membrane potential between WT and APPtg neurons: APPtg neurons were more depolarized (WT: -61.70±3.457 mV vs. APPtg: -58.67±6.427 mV; Fig. 3a). When we injected short step currents (3 ms current with 1 pA increment) CA1 pyramidal neurons in APPtg rats required lower current injections to initiate action potential firing (Short-pulse rheobase of 366.0 [263.3; 523.3] pA in APPtg vs. 503.0 [394.8; 664.5] pA in WT; Fig. 3b, c). Moreover, consistent with previous findings in mouse models^8^, CA1 pyramidal neurons of APPtg rats had an increased input resistance (by 20%) when compared to WT (92.98 [75.55; 129.2] MΩm; APPtg: 113.8 [86.22; 147.3] MΩm; Fig. 3d, e). Other electrophysiological parameters, such as sag ratio and firing frequency were not different between the two experimental groups (Fig. 3f). The observation that CA1 neurons require less depolarization to discharge is in agreement with our *in vivo* observation of neuronal hyperactivity. It suggests that an elevated intrinsic cellular excitability of CA1 pyramidal cells contributes to neuronal hyperactivity.

**Figure 3.**
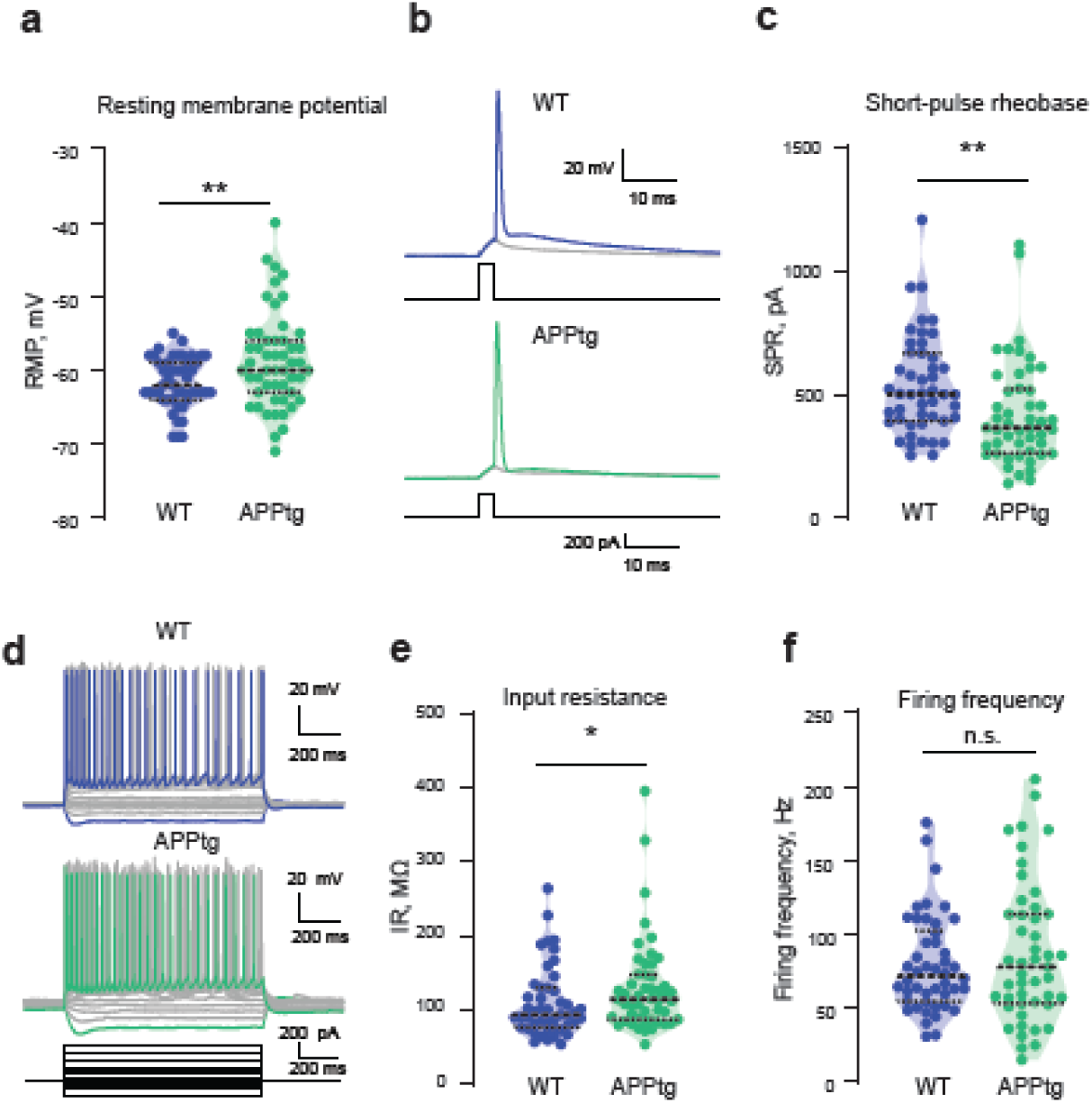
Increased intrinsic excitability of CA1 pyramidal neurons in APPtg rats. (a) APPtg neurons were more depolarized to comparison to WT ones. RMP reached -61.70 ± 3.46 in WT rats (n=47) and -58.67 ± 6.42 in APPtg rats (n=51) (unpaired t-test, p= 0.005) (b) Short pulse rheobase (SPR) detected by current injections with +1pA increment. Grey traces display the last subthreshold voltage response, and blue and green trace represents the first suprathreshold responses of WT and APPtg, respectively. The injected currents are depicted as black lines on the bottom. (c) SPR was reduced in APPtg (n=44) in comparison to WT (n=48) neurons (Mann-Whitney test, p=0.0015). (d) Representative examples of CA1 pyramidal neurons firing behavior in WT (blue traces: current injection of -200 and +500 pA) and APPtg (green traces: current injection of -200 and +500 pA). (e) Input resistance was increased in APPtg rats (Mann-Whitney test, p=0.039). (f) CA1 pyramidal neurons fired with initial frequency of 71.62 [54.59; 102.4] Hz in WT (n=46) and 77.76 [53.08; 113.6] APPtg (n=47) rats (Mann-Whitney test, p=0.65).

### Increased action potential duration in APPtg rats

We further investigated whether the active properties of CA1 pyramidal neurons differed between the experimental groups. We observed a prolongation of the action potential for short current injections or long current injections at the level of 50% of repolarization (Half- with short pulse rheobase (HWspr); WT: 0.81 [0.73; 0.94] ms vs. APPtg: 0.91 [0.84; 0.98] ms); Half width long current injection; WT: 0.83 [0.74; 1.06] ms vs. APPtg: 0.93 [0.86; 1.04] ms; Fig. 4a-d). Finally, we used multi electrode array (MEA) extracellular recordings to confirm our findings obtained in the patch clamp experiments on the population level (Fig 4 e-h). The particular advantage of MEA recordings is the undisturbed intracellular milieu of neurons during the recording. After recording from 60 electrodes in hippocampal slices, we performed spike-sorting to identify single units inside the multimodal signal. We compared the duration of the extracellular signal between WT and APPtg rats. Our results further support the findings of the whole-cell current-clamp recordings. The action potential duration was prolonged from 0.29 [0.21; 0.36] ms in WT rats to 0.31 [0.24; 0.38] ms in APPtg rats (Fig. 4h). Taken together, our data support the view that increased intrinsic neuronal excitability of CA1 pyramidal neurons is already present in the beginning of extracellular Aβ- deposition in APPtg rats.

**Figure 4.**
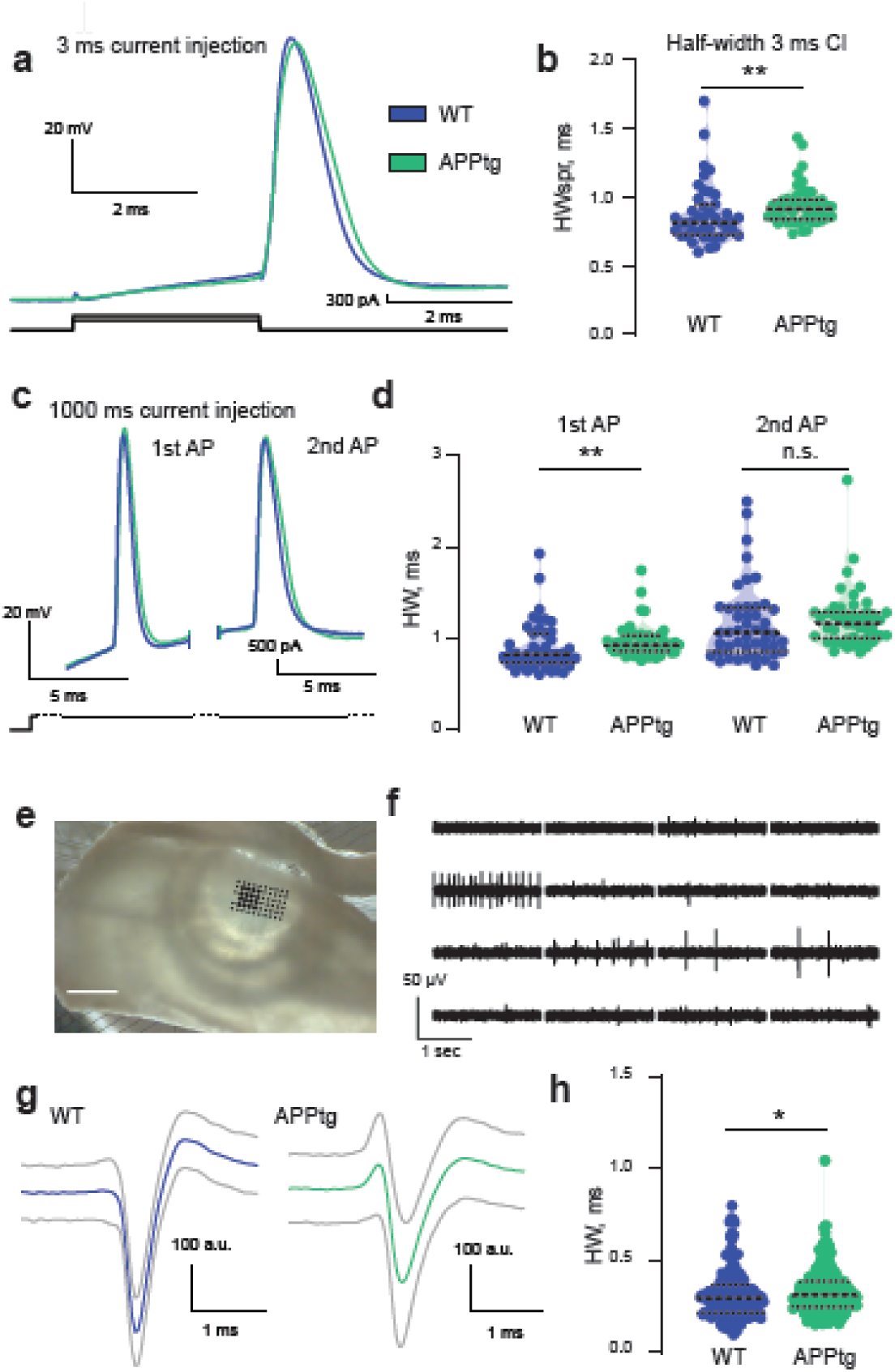
Increased action potential duration in APPtg rats. (a) Representative examples of single action potential in WT (blue) and APPtg (green) CA1 pyramidal neurons on the first suprathreshold responses following a 3ms current injection of 517pA and 352pA, respectively. (b) The half-width of the first suprathreshold action potentials was prolonged in APPtg rats in comparison to WT (n=44 for WT and n=48 for APPtg, Mann-Whitney test, p=0.002). (c) The first half-width (HW1) and second half-width (HW2) were determined as duration of the first and second action potential at the level of 50% repolarization for a +500 pA step current injection. (d) HW1 was prolonged (n=47 for WT and n=51 for APPtg, Mann- Whitney test, p= 0.001) and was not significantly different for the second action potential (Mann-Whitney test, p=0.087). (e-h) MEA recordings confirm prolongation of action potential duration in hippocampal slices of APPtg rats. (e) Hippocampal slice on a 6×10 electrodes MEA. Scale bar, 1 mm. (f) Original extracellular recordings, corresponding to the eight electrodes, depicted in bold black on the MEA slice image in e. (g) Examples of mean spike waveforms in WT (blue) and APPtg (green) rats. Grey lines correspond to SD values. (h) Distribution of extracellular spike half-widths in WT (199 units in 17 slices, 5 animals) and APPtg (194 units in 21 slices, 9 animals) rats. Extracellular measured action potentials were broadened in APPtg CA1 hippocampal neurons in comparison to WT ones (Mann-Whitney test, p=0.035).

## Discussion

Neuronal network dysfunction is thought to contribute to cognitive deficits in AD and under AD-like conditions in mouse models^9, 15^. Here we use a transgenic rat model to analyze hippocampal neuronal network function under AD-like conditions. Performing Ca^2+^-imaging in the hippocampus of anesthetized McGill-R-Thy1-APP rats *in vivo* revealed the presence of hyperactive CA1 neurons indicating changes on the neuronal microcircuit and network level. Although impaired inhibition has been previously shown to contribute to neuronal network dysfunction in mouse models^10, 21^, we did not detect inhibitory deficits in the measured parameters in the rat model. Instead, we found changes in passive electrical properties and intrinsic excitability that resulted in neuronal hyperexcitability. The changes in passive properties and intrinsic excitability observed in the present study suggest that intrinsic mechanisms precede circuit-based mechanisms during the progression of amyloidosis.

### Disturbance of the hippocampal neuronal network

Monitoring neuronal Ca^2+^-transients in the cortex of rats has been previously achieved using head-mounted microscopes^35^ or by voluntary head restrain in combination with a two-photon microscope^36^. However, so far two-photon *in vivo* imaging of deeper brain structures like the hippocampus has only been reported in mice^29, 37, 38, 39^. In this study we now have designed and used a specialized hippocampal cranial window that enabled us to perform hippocampal *in vivo* Ca^2+^-imaging in the hippocampus of rats. Two-photon imaging of Ca^2+^-transients in the cortex and hippocampus of mouse models with AD-like pathology has revealed hyperactive neurons in spatial proximity to Aβ-plaques^2^. Neuronal hyperactivity was observed both at pre-plaque stages and following Aβ-plaque deposition^16^. Using the genetically encoded Ca^2+^-indicator GCaMP6m, we identified hyperactive neurons in APPtg rats. Compared to the previous experiments performed in mice, overall neuronal activity in the hippocampus was lower. Apart from possible general activity differences between the two species within hippocampal networks, differences could be explained by the use of a different Ca^2+^-indicators, with largely different kinetics and sensitivities, but also difference in the way we detected Ca^2+^-transients (see Methods). In mice, treatment with a γ-secretase blocker to reduce the amount of Aβ-generation decreased the number of hyperactive neurons underscoring the importance of Aβ for neuronal hyperactivity^16^. Aberrant Ca^2+^- transient frequency was also detected in the visual cortex of awake mice with AD-like pathology during resting states^17^. Our *in vivo* Ca^2+^-imaging data confirm these findings and suggest that neuronal hyperactivity is species independent. It is possible, that similar pathomechanisms could be observed in AD patients, which exhibit signs of neuronal hyperactivity in depth recordings of the temporal lobe^40^.

### Intact inhibitory parameters at an early stage of Aβ-plaque deposition

Reduced inhibition represents one of the two current candidate mechanisms for increased neuronal excitability and hyperactivity^9, 15^. Impaired parvalbumin positive (PV^+^) GABAergic neurons, a subpopulation of GABAergic neurons involving fast-spiking, soma-targeting basket cells, but also several other cell types^41^, have been detected in mice with AD-like pathology. Interestingly, specific ectopic expression of Na^+^-channels in PV^+^ neurons with the rationale to increase PV interneuron activity was sufficient to improve memory in diseased mice^10^. Precisely timed optogenetic manipulation of hippocampal PV interneurons improved memory in mice with AD-like pathology^42, 43^. Furthermore, impaired innervation of hippocampal somatostatin positive GABAergic neurons contributed to reduced formation of postsynaptic spines and learning deficits in mice with AD-like pathology^21^. Somatostatin positive interneurons represent a population of GABAergic neurons comprising several cell types that predominantly targets apical dendrites of pyramidal neurons^10, 21^. Therefore, we tested whether impaired inhibition contributed to the observed hyperactive CA1 neurons in the rat model. Our protocol tested both for proximal and distal recurrent inhibition of CA1 pyramidal neurons in brain slices, as a shift of inhibition from proximal to distal dendritic regions has been reliably observed upon rhythmic and repeated stimulation in the theta and gamma range^33, 34, 44^. Our data revealed intact and unchanged IPSPs at early (more proximal inhibition) and delayed (more distal inhibition) stimulation time points (Fig. 2). Our findings suggest that other mechanisms than reduced inhibitory circuit function may underlie the neuronal hyperactivity that is observed in CA1 neurons at the beginning of Aβ-deposition.

### Mechanisms of increased excitability

In the present study, increased neuronal excitability has been shown for the first time under *in-vivo* and *in-vitro* experimental conditions in McGill-R-Thy1-APP rats at an early stage of AD-like pathology. We suppose that hyperexcitability may have multi-factorial reasons. Increased glutamatergic neurotransmission represents one hypothesis. Aβ for example binds to the ß2 adrenoreceptor, which through an intracellular cascade modulates glutamatergic transmission via AMPA receptors^45^. Furthermore, suppression of glutamate reuptake may be responsible for increased glutamatergic neurotransmission^46^. However, our findings support an intact glutamatergic neurotransmission in APPtg rats (Fig. 1j,k).

Another line of evidence follows reduced dendritic arborization, reduced cell size and loss of synapses that have been consistently detected in human AD patients^3, 47, 48, 49^ and in animal models^3, 4, 6, 7^. Modeling and experimental studies have shown that reduced cell size and membrane area may alter the input-output function of pyramidal cells by increasing the input resistance of neurons^8^. Such an increase in input resistance facilitates the firing of action potentials in response to synaptic input^8^. Indeed, APPtg rats exhibited an increased input resistance and reduced short-pulse rheobase at 6-9 months of age (Fig. 3). Such electrophysiological changes have not been observed at earlier stages (age 3-4 months) in the same APPtg rat model^23, 50^,. In addition, the increased action potential half-width observed in the present study further supports the view that intrinsic excitability changes of neurons contribute to neuronal hyperexcitability at early stages. Aß may affect large conductance calcium-activated potassium (BK) channels thereby contributing to the cellular hyperexcitability^51^. Mechanistically, a broadening of action potentials may facilitate and increase synaptic glutamate release^52^. Moreover, the synaptic input-output relationships were indistinguishable between 6–9 months old WT and APPtg rats, challenging the hypothesis of increased glutamatergic synaptic transmission.

Our findings suggest that early-stage neuronal hyperexcitability is mediated by changes in the intrinsic excitability of CA1 pyramidal neurons, rather than by synaptic changes. Thus, our data support the view that altered intrinsic excitability precedes inhibitory and excitatory synaptic dysfunction during disease progression.

## Methods

### Animals

Transgenic rats expressing human APP_751_ with Swedish (APP_KM670/671NL_) and Indiana (APP_V717V_) mutations under the control of the murine Thy1.2 promoter (McGill-R-Thy1-APP)^28^ of both genders were investigated. In this study we used 6-9 months old homozygous McGill-R-Thy1-APP rats (APPtg) and compared them with wildtype (WT) littermates. As previously published, homozygous APPtg rats displayed a behavioral phenotype as early as 3 months of age in the Morris water maze task^53, 54^ and in a visual discrimination task at the age of 4-6 months^55^. Furthermore, at the age of 7–10 months APPtg displayed altered profiles of social behavior and ultrasonic communication as well as increased locomotor activity during active periods of the circadian cycle^54^. To obtain homozygous APPtg and WT rats, we crossed heterozygous APPtg rats and identified homozygosity using a qPCR protocol. Furthermore, we carried out backcrossing with WT rats for one generation in some cases. McGill-R-Thy1- APP rats were genotyped as described previously^28, 53^. Briefly, genomic DNA was isolated from ear punches, using the peqGOLD Tissue DNA Mini Kit S-Line (PEQLAB Biotechnologie, Erlangen, Germany). The DNA concentration was measured by Peqlab NANODROP2000c (PEQLAB Biotechnologie, Erlangen, Germany). For distinguishing between homo- und heterozygous McGill-R-Thy1-APP rats qPCR was performed using the following primers: human APP-specific primers (forward: 5’ ATC CCA CTC GCA CAC AGC AG 3’, reverse: 5’ GGA ATC ACA AAG TGG GGA TG 3’) and GAPDH (forward: 5’ GGG GAA GGA CGC TGT ACG GG 3’, reverse 5’ AAG GGG AGC AAC AGC TGG GGT 3’). For qPCR each sample was processed 3 times in 2 dilutions with a total of 7ng and 14 ng of genomic DNA. The standard curve was determined on Applied Biosystems 7900HT Fast Real-Time PCR System (Applied Biosystems, Ca, USA) using 2 x PowerSYBR Green PCR Master Mix (Applied Biosystems, Ca, USA). The quantification was performed using 2^-ΔΔCt^ method and at least one sample of known heterozygous genotype served as internal standard. All experiments were performed in agreement with European Committees Council Directive (RL2010/63/EU) and were approved by the local authorities Landesamt für Natur, Umwelt und Verbraucherschutz (LANUV) of North Rhine-Westphalia.

### Slice preparation

Horizontal hippocampal slices were prepared as described previously^56^. In brief, rats were anesthetized with Ketamine (0.13 mg/g) and Xylazine (0.01 mg/g) before being decapitated. The brains were quickly transferred to ice cold sucrose solution, containing the following components in mM: 60 NaCl, 100 sucrose, 2.5 KCl, 1.25 NaH_2_PO_4_, 26 NaHCO_3_, 1 CaCl_2_, 5 MgCl_2_, 20 glucose, oxygenated with 95% O_2_ and 5% CO_2_. Horizontal hippocampal slices (300 *µ*m thickness) were prepared on a VT-1200S vibratome (Leica Microsystems, Wetzlar, Germany) and transferred to a submerged chamber at 35°C for 30 min and kept at room temperature thereafter in standard ACSF solution, containing in mM: 125 NaCl, 3 KCl, 1.25 NaH_2_PO_4_, 26 NaHCO_3_, 2.6 CaCl_2_, 1.3 MgCl_2_, 15 glucose.

### Whole cell patch-clamp recordings

Current-clamp recordings were performed using an BVC-700A amplifier (Dagan, Minneapolis, USA) with standard ACSF solution at 34°C containing the GABAB-receptor blocker CGP55845 (1 *µ*M, Tocris). Recording pipettes with a resistance of 3-5 MΩ were pulled with a DMZ universal electrode puller (Zeitz-Instruments Vertriebs-GmbH, Martinsried, Germany) and filled with intracellular solution of the following composition in mM: 140 K-gluconate, 7 KCl, 5 HEPES-acid, 0.5 MgCl_2_, 5 phosphocreatine, 0.16 EGTA. The liquid junction potential of -10 mV was not corrected. Signals were digitized at ≥25 kHz using an ITC-18 interface board controlled by custom-written procedures in Igor Pro 6.31. Resting membrane potential (RMP) was measured immediately after establishing the whole-cell configuration. Following that, the membrane potential was adjusted to −60 mV by continuous current injection. For estimation of the short-pulse rheobase (SPR), short 3 ms current pulses with increasing amplitude in steps of 1 pA were injected until reaching action potential threshold^8^. HWspr (half-width short-pulse rheobase) was calculated as the half-width of action potential (AP) duration at the level of 50% of repolarization of the first AP measured with the short pulse rheobase protocol using the TaroTools offline routines implemented in Igor Pro (6.31). Hyper- and depolarizing current steps of 1 s duration were applied to calculate the neuronal excitability Input resistance was calculated from linear fitting the range of current injections (−20, 0, 20 and 50 pA) with custom-made Igor Pro software (6.31). HW1 and HW2 represent the duration at half maximum amplitude of the first and second AP at the maximal current injection of +500 pA. Frequency was calculated as the instantaneous frequency between the first two APs following the same current injection. Sag ratio represents the ratio between steady membrane potential and peak potential resulting from a -200 pA current injection.

Recurrent inhibition was studied as described previously^33^. A cluster electrode (CE2F75, FHC, Bowdoin ME, USA) was placed onto the alveus close to the subiculum. Biphasic current pulses were generated using a stimulus isolator (A-M Systems, Model 2100, Sequim, WA, USA). To isolate CA1 originating recurrent inhibition, the subiculum as well as the CA3 subfield were cut off, sparing the alveus. Input-output curves were created using stimulus durations of 0.15–0.2 ms, stepwise increasing intensities by 0.025 mA up to 0.4 mA, using custom made online routine. Following that, 10 stimulations at theta frequency (5Hz) were applied, repeated 3 times and averaged. Additionally, 10 alveus stimulations in 100 Hz bursts at theta frequency (5 Hz) were performed^33^. Time-curve of IPSP changes were investigated and compared between WT and APPtg McGill rats. For 100 Hz bursts at theta frequency (5 Hz) the first IPSP component of the first pulse was compared with the first IPSP of the last pulse to estimate the contribution of more proximal inhibition (Fig. 2h). Accordingly, the third IPSP component of 100 Hz burst was compared to the third IPSP component of the last pulse to quantify more distal inhibition.

### Thioflavin-S staining and determination of plaque load

Acute horizontal hippocampal slices of 300 *µ*m thickness were fixed for 24 h in PFA (4%) and washed 2 times in 0.1 M PBS. The slices were preserved in 0.05% sodium azid in PBS at 4°C before staining. Slices were stained by bath application of 0.1% Thioflavin-S in deonized water for 20 minutes at RT. A total of 4 slices from 4 WT rats, 9 slices from APPtg rats (6-9 months), used for the patch-clamp experiments, and 3 slices from old (>18 months) APPtg rats used as a positive control, were stained.

Following the staining, slices were rinsed 3×10 min with PBS to remove the residual dye, Consecutive images through the hippocampus were made with a Zeiss LSM 700 confocal microscope from both sides of the slice using a 10× objective (Plan-NEOFLUAR 10×/0.3, Zeiss, Germany). The total number of Thioflavin-S positive plaques was determined in one hippocampal slice per rat and plaque density was calculated as amount of plaques per 1 mm^2^ of hippocampal area using ZEN black software.

### 6E10 and 4G8 staining and distinguishing between APP and Aß

Acute horizontal hippocampal slices of 300 *µ*m thickness were fixed for 24 h in PFA (4%) and washed 2 times in 0.1 M PBS. The slices were permeabilized in 0.5% Triton-X100 in PBS for 1 h. Then, the slices were incubated overnight in blocking reagent (10% normal goat serum, 1% Triton-X100 and 20% BSA) with primary antibodies – either anti-6E10 (1:1000, Covenance, mouse) and anti-NeuN (1:1000, Covenance, rabbit) or anti-4G8 (Covenance, mouse, 1:500) and anti-NeuN (1:1000,Covenance, rabbit). On the next day the slices were first washed 3 times with PBS. The slices were incubated for 1.5 h in 5% normal goat serum in PBS with secondary antibodies – either anti-mouse Alexa Fluor 488 (1:400, Life, goat,) and anti-rabbit Alexa Fluor 647 (1:400, Life, goat) or anti-mouse Alexa Fluor 405 (1:400, Life, goat) and Alexa Fluor 647 (1:400, Life, goat). The slices were washed 3 times with PBS and mounted with Dako mounting medium (Agilent). Fluorescence images of brain section were obtained using a confocal microscope (Zeiss LSM700).

### GFAP and Iba1 staining and determination of inflammatory response

Three APPtg and three WT rats at the age of 6–9 months and 1–3 months after hippocampal window surgery were transcardially perfused using sterile 0.1 M PBS and 4% PFA. The brain was removed and post-fixed in 4% PFA at 4°C overnight. For staining, free-floating coronal hippocampal slices of 50 *µ*m thickness were produced and incubated overnight in staining solution (4% normal goat serum, 0.4% Triton-X 100 and 4% BSA) with anti-GFAP antibody (1:1000, ThermoFisher, rat) and anti-Iba1 antibody (1:1000, Wako, rabbit). After washing 3 times with 0.1 M PBS, the free-floating sections were incubated for 2 h in staining solution together with anti-rat Alexa 594 and anti-rabbit Alexa 647 (1:400, ThermoFisher). Slices were washed 3 times with PBS and mounted using Dako Fluorescent mounting medium (Agilent). Fluorescence images of brain section were obtained using a confocal microscope (Zeiss LSM700).

### Microelectrode Array (MEA) recordings

For MEA recordings of spontaneous action potentials, hippocampal CA1 slices were prepared the same way as for patch-clamp recordings. After cutting, slices were transferred to an interface chamber (Brain Slice Chamber System with Haas Top, Warner Instruments, Hamden, USA) containing ACSF (Maier, Morris, Schmitz 2009) for recovery (mM): 119 NaCl, 2.5 KCl, 2.5 CaCl_2_, 1.0 NaH_2_PO_4_, 26 NaHCO_3_, 1.3 MgCl_2_, 10 glucose, oxygenated with 95% O_2_ and 5% CO_2_. Inside the interface chamber, hippocampal CA1 slices were kept on lens cleaning tissue (Grade 105, Whatman, Maidenstone, England) allowing optimal oxygenation due to a laminar flow of preheated (35 °C) ACSF above and underneath the slices for at least 3 hours. Extracellular waveforms were recorded with a MEA2100-System (Multi Channel Systems, Reutlingen, Germany) on 60pMEA100/30iR-Ti MEAs with round titanium nitride (TiN) electrodes. Electrodes were arranged in a 6×10 matrix, with a spacing of 100 *µ*m and an electrode diameter of 30 *µ*m. ACSF temperature was adjusted to 35 °C using a heatable perfusion cannula PH01 together with a TC01 controlling unit (Multi Channel Systems, Reutlingen, Germany). A constant negative pressure of 25-30mBar mbar was applied to the perforated MEAs to stabilize slice position, improve electrode connection, and maintain vertical ACSF flow. Data were sampled at 25 kHz with an MEA2100-lite-Interface Board and acquired with MC_Rack (V 4.5.16.0, Multi Channel Systems, Reutlingen, Germany).

Ten-minutes MEA recording epochs were analyzed using the open-access template based unit-detection and -sorting algorithm Kilosort) ^57^ (https://github.com/cortex-lab/KiloSort) in Matlab (MathWorks, R2016a). The following parameters were used for the spike sorting: ops.Nfilt = 256; ops.whitening = ‘full’; ops.nSkipCov = 1; ops.whiteningRange = 32; ops.criterionNoiseChannels = 0.2; ops.Nrank = 3; ops.nfullpasses = 6; ops.maxFR = 20000; ops.fshigh = 500; ops.fslow = 2000; ops.ntbuff = 64; ops.scaleproc = 200; ops.NT = 32*1024+ ops.ntbuff; ops.Th = [2 8 8]; ops.lam = [5 20 20]; ops.nannealpasses = 4; ops.momentum = 1./[20 400]; ops.shuffle_clusters = 1; ops.mergeT = .1; ops.splitT = .1; ops.initialize = ‘fromData’; ops.spkTh = -5; ops.loc_range = [3 1]; ops.long_range = [30 6]; ops.maskMaxChannels = 5; ops.crit = .65; ops.nFiltMax = 10000.

Automatically detected units were manually validated using Phi^58^. For an identified unit the mean waveform was calculated from 2000 random spikes events corresponding to this unit. Half-width of the extracellular spikes was determined at the half-maximum of the mean spike waveforms negative deflection.

### Data analysis

Analysis of calcium fluorescence signal was performed using custom analysis software routines in Fiji and Igor Pro. Motion correction of calcium time series was performed using the batch registration function of the custom Fiji macro TurboReg (Dr. Philippe Thévenaz, Swiss Federal Institute of Technology).

Cells were manually identified from the maximum intensity projection of single-trial time series and marked as region of interest (ROI) using Fiji. Change in calcium intensity over time was obtained for every ROI. Calcium signals were depicted as relative fluorescence change, ΔF/F. The ΔF/F was calculated by subtracting the baseline fluorescence F0 from the signal and dividing the value by F0, ΔF/F = (F – F0)/F0. F0 was defined as the mean of the smallest 40% of all values in a time series. For further analysis we used the custom macro Taro Tools (Dr. Taro Ishikawa, https://sites.google.com/site/tarotoolsregister/) for Igor Pro 6. The ΔF/F traces were smoothed using a 5-point, box-smoothing filter. Calcium peaks were identified according to following criteria: First of all, a baseline ΔF/F for every peak was calculated as the mean of the 5s-window preceding the peak of the event. Calcium events were counted when the peak was three times the SD value of the baseline. Second, the ΔF/F amplitude of the smoothed calcium event had to be at least 10% since this is the value that has been described as the calcium increase in response to one action potential in the visual cortex^59^. The third precondition was that two consecutive events had to be separated by at least 500 ms.

Classification of neurons was performed according to previous reports by Busche *et al*. ^2^. The first category, termed ‘silent cells’, comprises neurons that show less than 0.4 Ca^2+^- transients/min. The second category, termed ‘normal cells’, represents cells that displayed between 0.4 and 4 Ca^2+^-transients/min. The third category, termed ‘hyperactive cells, includes all neurons with more than 4 Ca^2+^-transients/min.

Electrophysiological data were recorded and analyzed using Igor Pro (WaveMetrics, Lake Oswego, Oregon, USA). Statistical comparisons were performed with Prism 7 and 8 for Windows (Prism version 7.0.3 and 8.0.2 for Windows, GraphPad Software, La Jolla California USA).

Evoked IPSP were fitted using the function f(t) = f_0_ - A*(1-exp(-(t-t_0_)/□_r_))^^5^*exp(-(t-t_0_)/□_d_) ^60^, where f_0_ is a baseline, A is a constant □_r_ and □_d_ are the rise and decay time constants. IPSP rise and decay time constants were averaged from 3 repetitive measurements and compared between WT and APPtg.

### Virus injection

To conditionally express the fluorescent calcium indicator GCaMP6m^59^ in CA1 pyramidal neurons, we anesthetized female and male rats by injecting - either intraperitoneally (i.p.) or intramuscularly (i.m.) - a mixture of Ketamine/Xylazine (0.13/0.01 mg/g). When the animal was asleep, we made a small incision using a scalpel and performed a small craniotomy (Ø 1 mm) using a dental drill at a previously marked spot overlying the dorsal hippocampus at the coordinates -4.2 mm anterior posterior and +2.6 mm mediolateral from bregma. We then injected three times 1 *µ*L (100 nL/min) of *AAV2-hSyn-GCaMP6m-WPRE* at each 3.3 mm, 3 mm and 2.7 mm dorsoventral from the skull surface using a microliter syringe (Hamilton, USA) controlled by a four-channel micro controller (SYS-Micro4, World Precision Instruments, USA). After the last injection step, we kept the needle in place for 10 min so that the virus had time to diffuse into the tissue. The needle was carefully removed and the skin was stitched.

### Hippocampal window preparation

The preparation of the cranial window was carried out similar to previously described procedures in mice^38^, including some refinements. The animals were anesthetized by injecting - either i.p. or i.m. - a mixture of Ketamine/Xylazine (0.13/0.01 mg/g). During the procedure the body temperature was maintained at ∼37 °C using a heating plate. The eyes of the rat were covered with eye ointment (Bepanthen, Bayer, Germany) during the surgery. After hair removal with tweezers and disinfecting with 70% ethanol, the skin was opened with a scalpel. The exposed cranial bone was cleaned and dried with Sugi absorbent swabs (Kettenbach GmbH, Germany). The bone surface was scraped and cleaned with a scalpel. For preparing a base-layer of glue to which subsequently all parts of the headholding set were fixed, we used a two-component dental glue (OptiBond FL, Kerr, USA). After drying and roughening of the skull, a thin layer of the priming component was applied for 15 seconds to the skull surface with a slight scrubbing motion. The layer was gently air-dried for 5 seconds. Using the same applicator-brush, a layer of the adhesive component was applied to the skull surface the same way as it was done with the priming component. The glue was then light cured for 20 seconds. Using a dental drill, we performed a craniotomy (Ø 5 mm) around the injection site. Cortical tissue was gently aspirated with a blunt 27 gauge needle until the external capsule became visible. A stainless steel cannula (Ø 5 mm, 2.5 mm height) covered with a cover glass (Ø 5 mm, 0.17 mm thickness) was inserted into the free space between hippocampus and skull. The small cavity between skull and cannula was sealed by with a mixture of dental acrylic cement (Cyano Veneer, Hagerwerken, Germany) and dental acrylic glue (Cyano Retarder, Hagerwerken, Germany). Using dental cement, a metal headpiece was glued to the skull on the contralateral side in order to allow fixation of the animal under the microscope. After surgery, the animals were treated with Carprofen (5 mg/kg) two times per day for two days and treated with antibiotics.

### Two-photon calcium imaging

For two-photon imaging we used a TriM Scope II (LaVision BioTec GmbH, Germany) microscope and a 25x 4mm-working-distance, water-immersion objective (numerical aperture = 1, model XLPLN25XSVMP2, Olympus). Imaging was performed after ceasing of surgery-induced inflammation response, which in mice takes four weeks^29^. For excitation of GCaMP6m at 920 nm we used a standard titanium sapphire (Ti:Sa) laser (∼900 fs laser pulse width at 80 MHz; Coherent). Fluorescence was assessed using a bandpass filter (510/20 nm; Semrock). For imaging under anesthesia, the rats were anesthetized with 1% isoflurane in oxygen using an isoflurane vaporizer (Kent Scientific, USA) and fixed with the metal bar to a holder in order to allow stable imaging.

### Electrophysiological recording in freely behaving rats

Recording was carried out in chronically implanted freely behaving 6–9 months old rats. The implantation procedure was carried out under recovery anaesthesia using a mixture of ketamine (80 mg/kg) and xylazine (8 mg/kg) (both i.p.) according to methods similar to those described previously^23^. Field EPSPs were recorded from the stratum radiatum in the CA1 area of the dorsal hippocampus in response to stimulation of the ipsilateral Schaffer collateral/commissural pathway. Monopolar recording electrodes (75 μm inner core diameter, 112 μm external diameter) and twisted bipolar stimulating electrodes (50 μm inner core diameter, 75 μm external diameter) were constructed from Teflon coated tungsten wires. The recording site was located 3.8 mm posterior to bregma and 2.5 mm lateral to midline, and the stimulating site was located 4.6 mm posterior to bregma and 3.8 mm lateral to midline. The final placement of electrodes was optimized by using electrophysiological criteria and confirmed via post-mortem analysis. Animals were allowed recover for at least 2 weeks after implantation surgery before recording from them as they freely explored a recording box in a well-lit room. The recording box consisted of the base of the home cage, including normal bedding and food/water, with the sides replaced with translucent Perspex walls (27 x 22 x 30 cm) and an open roof. The rats had access to food and water throughout the whole recording session from the same position as in the home cage. All animals were first habituated to the recording procedure over the post-surgery recovery period. Test EPSPs were evoked by square wave pulses (0.2 ms duration) at a frequency of 0.033 Hz and an intensity that triggered a 50% maximum response as determined after constructing an input-output curve.

## Statistical information

Shapiro-Wilko test was performed to test for normality for data sets with n<50, Kolmogorov-Smirnov test was performed to test for normality for data sets with n>50. Unpaired student’s t-test analysis (two-tailed), Mann-Whitney test or Kolmogorov-Smirnov test were performed where applicable. In experiments where more conditions were evaluated, one-way or two-way RM ANOVAs were performed followed by multiple comparisons testing using the Sidak’s correction method. * p<0.05, ** p<0.01, *** p<0.001.

## Data availability

The datasets generated during and/or analyzed during the current study are available within the article or the supplementary information files or from the corresponding authors upon reasonable request.

## Acknowledgements

This work was supported by CoEN/2011/11, the Deutsche Forschungsgemeinschaft (DFG, German Research Foundation – SFB 1089 C01, B06, C05 to MF and SR) and by Science Foundation Ireland, grant no. 14/IA/2571. We are grateful for the expert technical assistance of Meltem Eryilmaz.

## Author contributions

M.F., S.R., A.C.C., M.R., L.S. and M.M. were involved in the design of the study. A.C.C. participated in initial research planning discussions and the provision of well characterized McGill-R-Thy1-APP transgenic breeders. M.F., S.R., L.S., M.M. and H.-R.G. co-wrote the paper. L.S., Y.Q. and H.-R.G. performed the electrophysiology experiments and analyzed the data and prepared figures. I.K. assisted in designing electrophysiology recordings. M.M. and K.H. conducted the Ca^2+^-imaging. M.M. analyzed the Ca^2+^-imaging data and prepared figures. N.H. did morphological reconstructions. K.K. and F.F. provided microscopy assistance. J.S., D.F. did the histological stainings and prepared figures. D.J. provided data analysis assistance.

## Material & Correspondence

Materials are available on request. All correspondence should be directed to Martin Fuhrmann or Stefan Remy.

## Supplementary information

**Figure S1.**
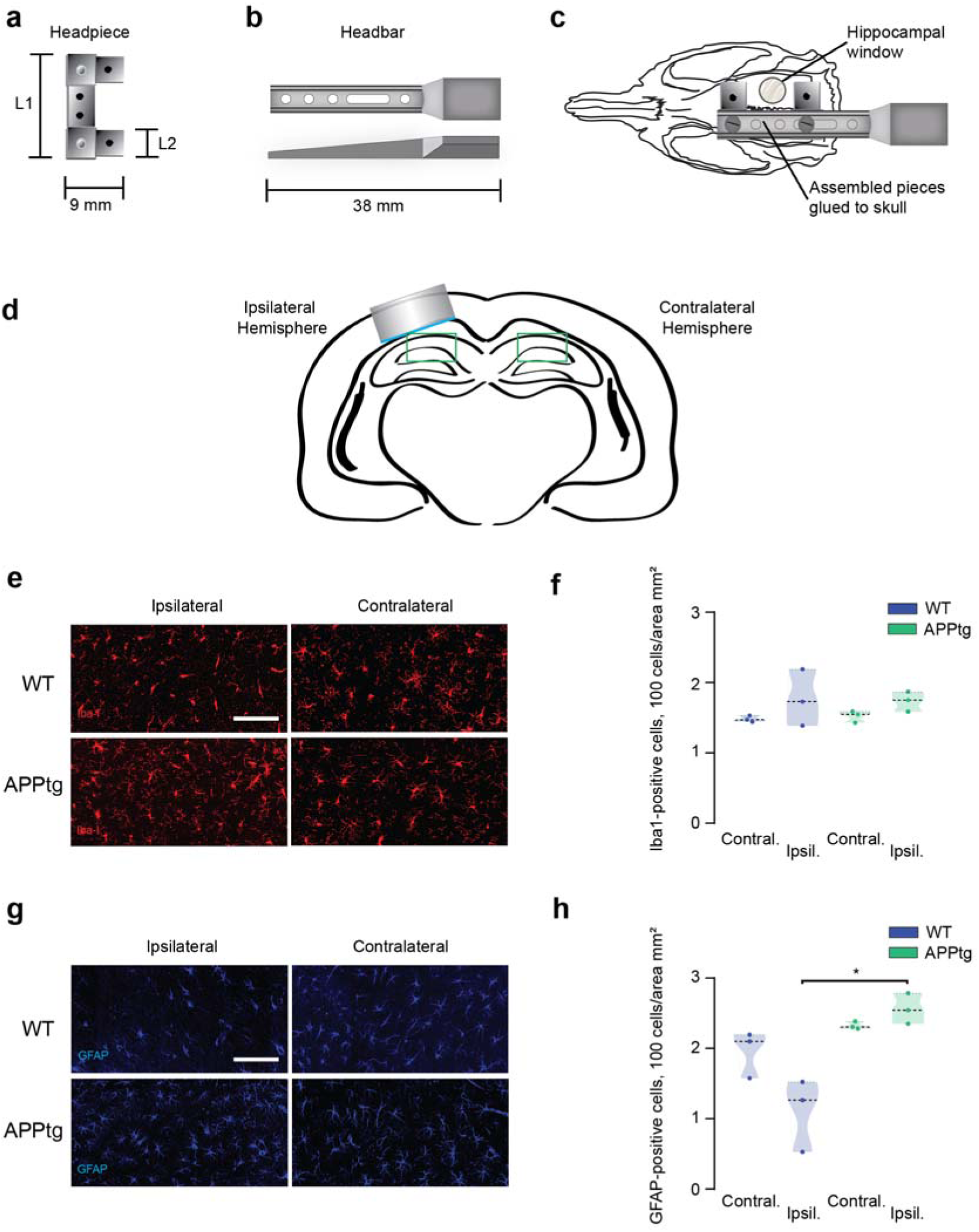
(a) Schematic illustration of the headpiece that was glued to the skull of the rat. Depending on the size of the head of the rats, headpieces with different length parameters (L1 and L2) were used. (b) Schematic illustration of head bar that was screwed to the headpiece. (c) Schematic illustration of assembled pieces fixed to the head of the rat. (d) Schematic illustration of hippocampal window implantation. (e) Comparison of immunohistochemical staining of microglia (red, Iba1) of ipsilateral (left) and contralateral (right) hemisphere in WT rats (top) and APPtg rats (bottom). (f) Quantitative analysis of Iba1- positive cells on WT contralateral (n=3, 1.47 [1.44; 1.53]) and ipsilateral (n=3, 1.73 [1.39; 1.53]) hemisphere as well as APPtg contralateral (n=3, 1.55 [1.43; 1.59]) and ipsilateral (n=3, 1.75 [1.59; 1.87]) hemisphere (Kruskal-Wallis Test, p>0.05). (g) Comparison of immunohistochemical staining of astrocytes (blue, GFAP) of ipsilateral (left) and contralateral (right) hemisphere in WT rats (top) and APPtg rats (bottom). (h) Quantitative analysis of GFAP-positive cells on WT contralateral (n=3, 2.1 [1.58; 2.19]) and ipsilateral (n=3, 1.27 [0.53; 1.52]) hemisphere as well as APPtg contralateral (n=3, 2.3 [2.28; 2.38]) and ipsilateral (n=3, 2.54 [2.35; 2.79]) hemisphere (Kruskal-Wallis Test, p=0.019). Scale bar=50 *µ*m

**Figure S2.**
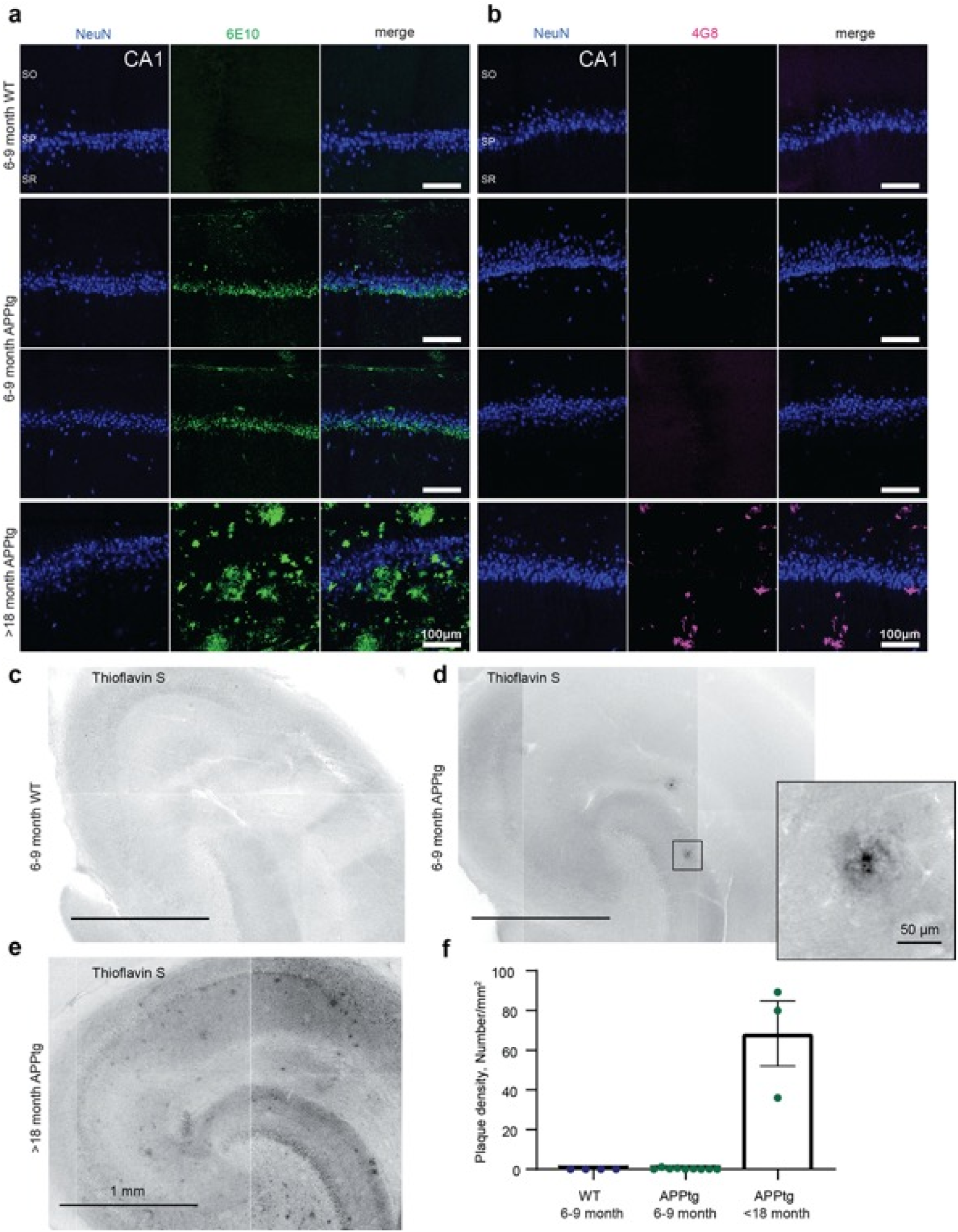
(a) NeuN and 6E10 staining of CA1 hippocampal region of 4 exemplary brains (WT 6-9 months, APPtg 6-9 months, APPtg >18 months). Scale bar 100 *µ*m. (b) NeuN and 4G8 staining of CA1 hippocampal region of 4 exemplary brains (WT 6-9 months, APPtg 6-9 months, APPtg >18 months). SO - stratum oriens, SP - stratum pyramidale, SR - stratum radiatum. Scale bar 100 *µ*m. (c), (d), (e) Thioflavin S staining of hippocampal region of 3 exemplary brains. Scale bar 1 mm. Inset scale bar 50 *µ*m. (f) Plaque density calculated for 1 mm^2^ in 3 groups of APPtg rats: WT (n=4), APPtg 6-9 months (n=9) and APPtg >18 months (n=3).

**Figure S3.**
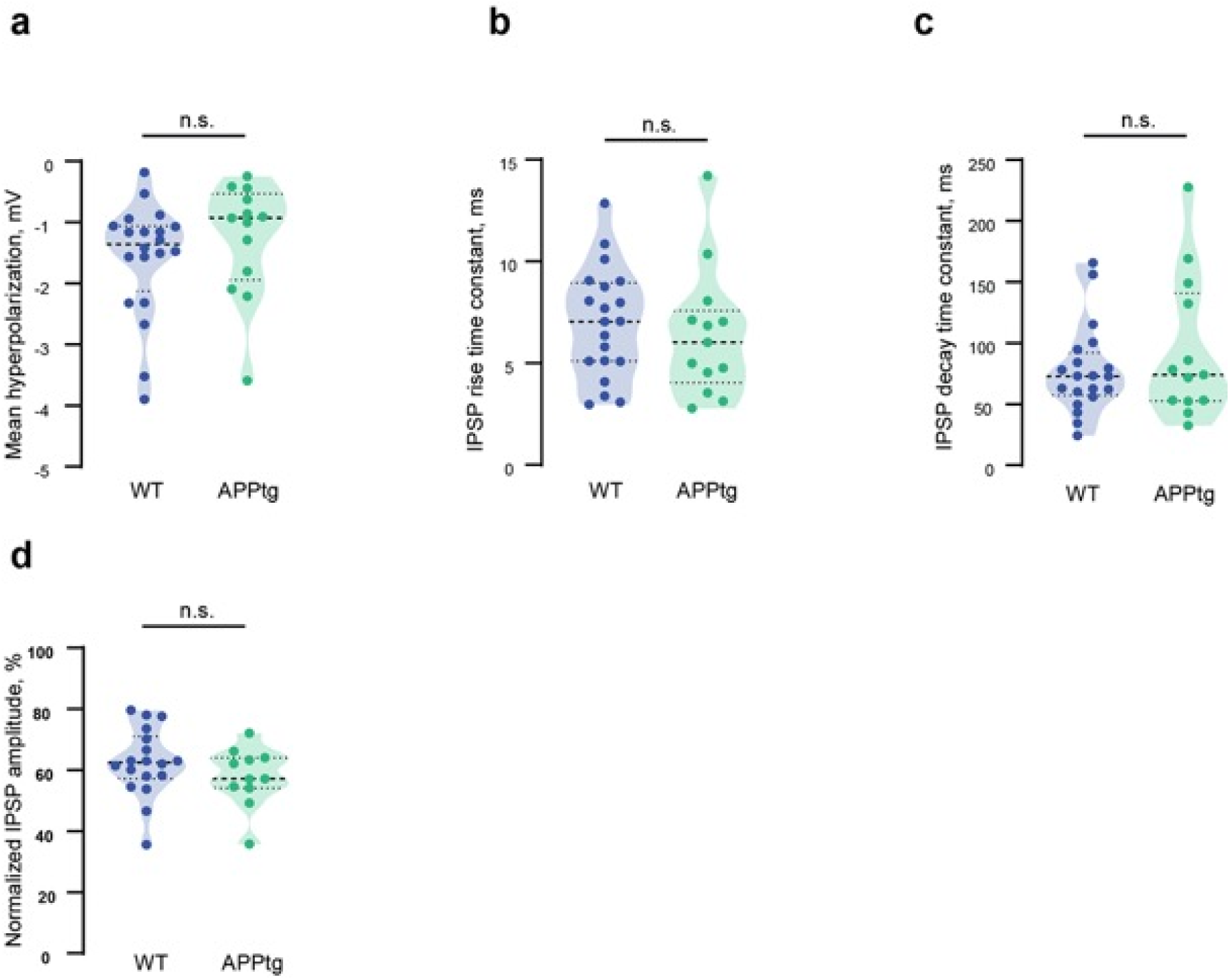
IPSP characteristics in CA1 pyramidal neurons evoked by alveus stimulation. (a) Mean membrane potential hyperpolarization was not different between WT (n=20 in 12 animals) and APPtg rats (n=13 in 7 animals) rats (Mann-Whitney test, p=0.158). (b,c) Time constants did not deviate between WT (n=20 in 12 animals) and APPtg rats (n=13 in 7 animals) rats: (b) IPSP time rise constant (Mann-Whitney test, p=0.392), (c) IPSP time decay constant (Mann-Whitney test, p=0.73). (d) Reduction in compound IPSP amplitudes 10th (last) vs. first stimulation (100 Hz burst at 5 Hz), was not different between WT (n=18 in 12 animals) and APPtg (n=11 in 6 animals) rats (unpaired t-test, p=0.259)

